# Value-based search efficiency is encoded in substantia nigra reticulata firing rate, spiking irregularity and local field potential

**DOI:** 10.1101/2023.05.28.542612

**Authors:** Abdolvahed Narmashiri, Mojtaba Abbaszadeh, Mohammad Hossein Nadian, Ali Ghazizadeh

**Affiliations:** Bio-intelligence Research Unit, Sharif Brain Center, Electrical Engineering Department, Sharif University of Technology, Tehran, Iran; School of Cognitive Sciences, Institute for Research in Fundamental Sciences (IPM), Tehran, Iran

**Keywords:** value memory, visual search, search efficiency, single-unit recording, basal ganglia, substantia nigra reticulata, macaque monkey, LFP, spiking variability

## Abstract

Recent results show that valuable objects can pop-out in visual search yet its neural mechanisms remain unexplored. Given the role of substantia nigra reticulata (SNr) in object value memory and control of gaze, we recorded its single unit activity while male macaque monkeys engaged in efficient or inefficient search for a valuable target object among low-value objects. Results showed that efficient search was concurrent with stronger inhibition and higher spiking irregularity in target present (TP) compared to target absent (TA) trials in SNr. Importantly, the firing rate differentiation of TP and TA trials happened within ∼100ms of display onset and its magnitude was significantly correlated with the search times and slopes (aka search efficiency). Time-frequency analyses of local field potential (LFP) after display onset revealed significant modulations of gamma band power with search efficiency. The greater reduction of SNr firing in TP trials in efficient search can create a stronger disinhibition of downstream superior colliculus which in turn can facilitate saccade to obtain valuable targets in competitive environments.

**Significant statement:** Most times we have to find a few relevant or highly valued objects among many objects that surround us. When our target objects are not distinct from their surroundings based on low-level features, searching for them becomes tedious and slow. Nevertheless, recent findings show that valuable objects can be found efficiently and fast if they have been repeatedly paired with reward. Our results show that the rate and pattern of spiking as well as local voltage fluctuations in the basal ganglia output which is known to control attention and saccade correlate with such value-driven search efficiency. Thus, in addition to reward learning, basal ganglia can have a role in skillful interactions with and rapid detection of rewarding objects.

## Introduction

Primates, including humans and monkeys, often face the challenge of quickly identifying valuable objects amongst many irrelevant or low-value objects in their surroundings. It has been long thought that visual search for targets that are not distinct from their surrounding by low-level guiding features such as color, size, or orientation should be invariably a slow and serial process requiring multiple shifts of gaze(Wolfe, 2021). Contrary to this classical belief, we have shown that object value association especially when over-trained, can lead to efficient value-based search with minimal dependance on set size, a phenomenon we refer to as ‘value-popout’ (Ghazizadeh et al., 2016). However, while the efficient visual search based on low-level guiding features can be easily explained by the parallel processing of color and orientation across the visual field, the neural mechanism that allows for efficient value-based search is currently unknown.

The neural circuitry underlying object value memories has been elucidated by single-cell recording studies, implicating key regions within the basal ganglia (BG), such as the ventral putamen (vPut) (Kunimatsu et al., 2019), caudoventral globus pallidus (cvGPe) (Kim et al., 2017), caudate tail (CDt) (Kim and Hikosaka, 2013; Yamamoto et al., 2013), and substantia nigra reticulata (SNr) (Yasuda et al., 2012; Ghazizadeh and Hikosaka, 2021, 2022a). In particular, the strongest discrimination between high and low-value objects is found in SNr neurons compared to any other region BG or even the cortex (Hikosaka et al., 2014; Ghazizadeh and Hikosaka, 2021). SNr neurons exert powerful inhibitory influence on saccades by projecting to the superior colliculus (SC) and are capable of rapidly distinguishing between objects with high and low values, within the first 100-150ms of object onsets (Yasuda et al., 2012). When faced with a single valuable object, SNr neurons reduce their firing to disinhibit saccade and when encountering low value object SNr neurons increase their firing to inhibit saccades. Nevertheless, the role of SNr in visual search when multiple objects are simultaneously present and its role in enabling efficient value-based search is not previously studied.

To address this issue, we recorded single-unit activity in SNr in monkeys engaged in value-based search task with low or high search efficiency. Value-based search task consisted of trials with a single high value object among low-value objects (target-present or TP trials) and trials in which all objects where low value (target absent or TA trials). We looked at the effect of trial type and search efficiency in the first and second order statistics of firing rate as well as on local field potential power across multiple frequency bands in SNr. Briefly, results revealed that there was a stronger discrimination of TP and TA trials based on SNr firing rate and firing irregularity during efficient search. Search efficiency was also reflected and correlated firing rate changes and gamma band modulation in LFP within SNr. These findings implicate BG circuitry and in particular its output emanating from SNr in value-popout phenomenon.

## Material and Methods

### Subjects and surgery

In our experiments, we utilized two rhesus macaque monkeys (male), monkeys J and P, who were 8 and 7 years old, respectively. The experiment was conducted in accordance with ethical standards for animal care and use established by the National Institutes of Health (NIH) and after obtaining approval from the local ethics committee at the Institute for Research in Fundamental Sciences (IPM) (protocol number 99/60/1/172). Prior to the experiments, each monkey underwent sterile surgery, during which a head holder and recording chamber were implanted on their heads while they were under general anesthesia. A rectangular chamber was positioned laterally over the right hemisphere of each monkey. In a subsequent surgery, craniotomies for SNr were conducted after the monkeys learned the experimental tasks on the right hemisphere of both monkeys, following MRI confirmation of the recording chamber’s location. The recording was performed using grids that had a spacing of 1 millimeter and were positioned on top of the chamber.

### Recording localization

To pinpoint the location of the SNr in both monkeys, T1-and T2-weighted MRI (3 T, Prisma Siemens) were employed. T2-weighted MRI was found to be particularly beneficial in imaging the SNr region due to its higher iron concentration (Yasuda et al., 2012). Additionally, the recording chambers used for both monkeys were filled with gadolinium to enhance contrast during imaging. The standard monkey atlas (D99 atlas; Reveley et al., 2017) was transferred into each monkey’s native spaces with the use of the MATres software (Nadian et al., 2023) to further confirm the location of SNr in each monkey. Consequently, we ascertained the region of the SNr that was reachable for recording in each monkey via the recording chamber (Fig 1G).

**Fig 1.**
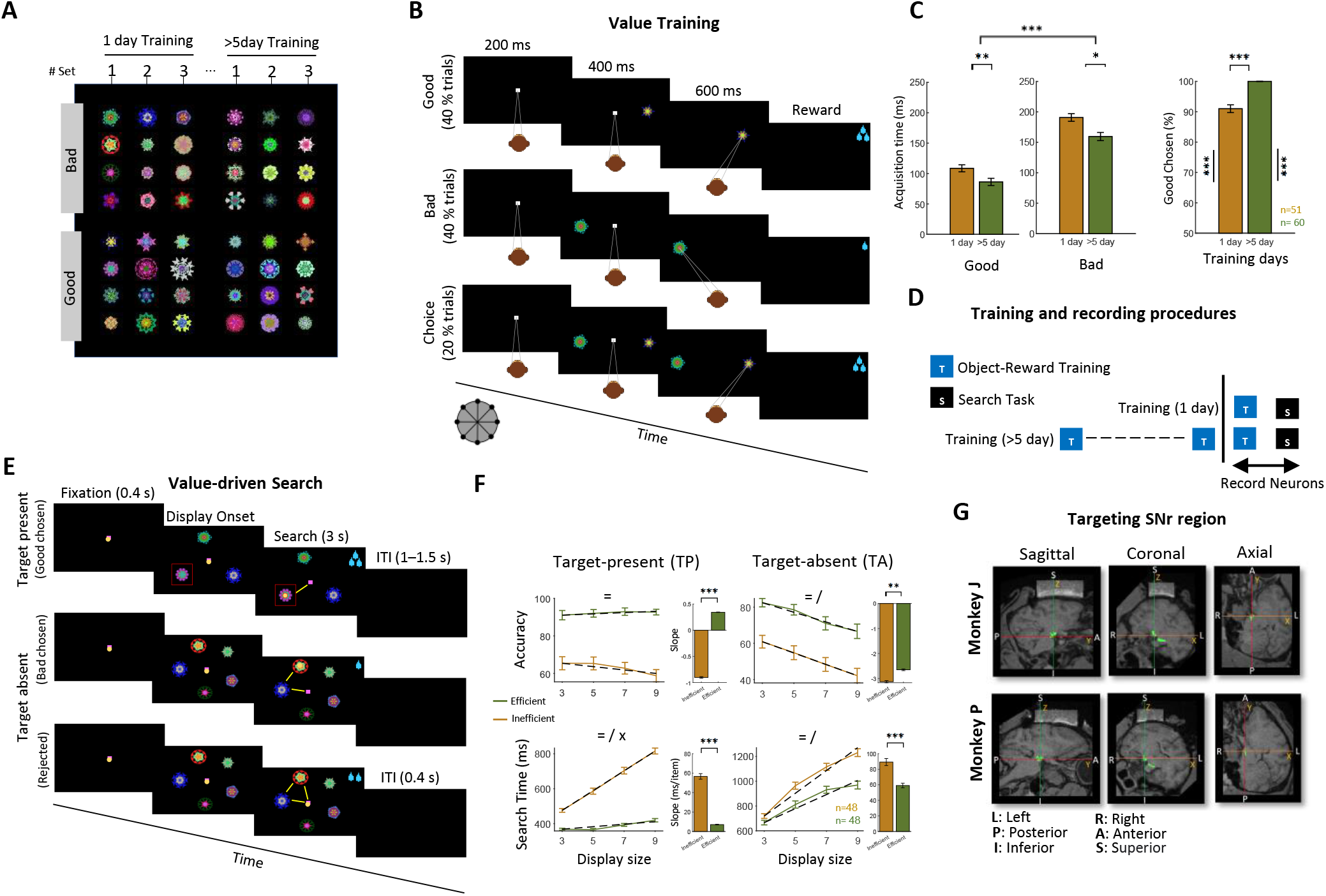
Value-based search paradigm and stimuli. A) The monkeys were trained with >400 fractal objects, which were divided into sets of eight, consisting of four fractals with high reward and four with low reward associations. These sets of fractals were subjected to value-based training, which lasted either for 1 day or >5 days. B) During the value training phase, the subjects participated in two types of trials: forced and choice trials which were intermixed. In forced trials, a single object (either good or bad) appeared randomly in one of 8 locations on the screen after a central fixation point, and the subjects had to make a saccade toward the presented object to receive a reward. In choice trials, two fractals (one good and one bad) appeared randomly in two of the 8 locations on the screen after a central fixation point, and the subjects had to choose one of the presented fractals to deliver the reward. C) The performance of the monkeys in the value training task. The time taken to acquire good objects and bad objects in forced trials (left two plots). The percentage of good object choice in choice trials (right plot) separately for objects that were trained for 1 day and objects that were trained for >5 days. D) Search task is done using distinct sets of reward-biased fractals with varying training durations (1-day and >5days). E) The search task consisted of target-present (TP) and target-absent (TA) trials. In each trial, a varying number of fractals (3, 5, 7, or 9) were displayed at equal distance on a circle 9.5° eccentricity. In TP trials, only one of the displayed fractals was good, while in TA trials, all of the displayed fractals were bad. The monkey could get large reward after finding and fixating the good object in TP trials and could proceed rapidly to the next trial by rejecting TA trials. F) Percentage of trials finding the good objects in TP trials and in rejecting the trial in TA trials (accuracy, top row) and search time taken to find good objects in TP or reject the trial in TA trials (bottom row), for both efficient and inefficient. The dotted line represents the linear fit and the inset bar plots indicate slopes. G) Sagittal, coronal, and axial views of the SNr region of two monkey brains that have been registered standard brain atlas (D99) for monkey J (top) and monkey P (bottom). The SNr regions are shown in green, and the recording chambers filled with gadolinium are shown in white. The MRI coordinates were determined as follows: X-axis represents the direction from left to right, Y-axis represents the direction from posterior to anterior, and Z-axis represents the direction from inferior to superior. The symbols =, /, and x represent the main effects of search efficiency, display size, and their interaction, respectively. The symbols *, **, and *** indicate statistical significance levels at p < 0.05, p < 0.01, and p < 0.001, respectively here and thereafter.

### Stimuli

Fractal-shaped objects were utilized as visual stimuli (Miyashita et al., 1991), as illustrated in Fig 1A. The fractals used in the experiment consisted of four polygons that were symmetrically arranged around a central core. Each polygon had smaller polygons in front of it, and their properties such as size, edges, and color were randomly selected. On average, the diameter of the fractals subtended 4 degrees in this experiment and shown on a CRT monitor. The monkeys were exposed to a substantial number of fractals, totaling over 400, organized in sets of eight. Each set was either trained in one reward training session or in more than five reward training sessions (over-trained) and were used in visual search task subsequently.

### Task control and neural recording

A customized software program written in C# (C-sharp) was used to present the task and record behavioral events and communicate with the neural data acquisition system (Cerebus Blackrock Microsystems Inc., Salt Lake City, UT, USA). Eye position was tracked at a rate of 1 kHz using an Eyelink 1000 Plus system (Ontario, Canada). Rewards were delivered in the form of apple juice diluted with water (at a 50% concentration for both monkeys), with varying amounts ranging from small (0.08 mL for both monkeys) to large (0.21 mL for both monkeys). During the experiment, the headfixed monkeys sat in a restraint chair and were presented with visual stimuli on a 21-inch CRT monitor during each experimental session.

Single-unit activity was recorded using tungsten electrodes coated with epoxy (FHC, 200 μm thickness). To prepare for each recording session, a sharp stainless steel guide tube was used to hold the electrode. The guide tube was then utilized to puncture the dura, allowing for the electrode to be cautiously inserted into the brain with the aid of an oil-driven Narishige (MO-97A, Japan) micromanipulator.

The electrical signal from the electrode was converted to a digital signal with a sampling rate of 30 kHz. Subsequently, the signals were amplified and filtered to isolate frequencies ranging from 1 Hz to 10 kHz and Blackrock’s online sorting was used for online survey of available single or multi-units. The resulting spike time data for the identified unit were then transmitted to MATLAB for generating online raster plots and peristimulus time histograms (PSTH). The neural recording data was processed using the Plexon offline sorter (Plexon, Dallas, TX, USA) in order to detect and organize the spikes. Standard spike sorting procedures were employed, including the application of a K-means supervised clustering algorithm, principle-component analysis (PCA), waveform template matching, and inter-spike interval distribution (ISI) analysis, among others. Initially, a voltage threshold was established for single channel, and spikes that surpassed this threshold were selected for further spike sorting analysis. The K-means clustering algorithm and manually were then utilized to separate different types of neurons, based on the observation of clusters formed by PCA in a 3-dimensional (3D) space, as well as the calculation of inter-spike interval (ISI) histograms. This study only considered neurons that were well-isolated and exhibited responsiveness to visual stimuli. A total of 50 neurons were recorded during both efficient or inefficient searches, comprising 28 neurons from monkey J and 22 from monkey P. To ensure consistency in our analysis, we applied the search slope criteria, as described in the results section. As a result, we utilized 48 neurons (27 from monkey J and 21 from monkey P) for further analysis, out of which 39 neurons were recorded during both types of searches (21 from monkey J and 18 from monkey P).

### Neural data analysis

All neural spiking data were aligned with the visual display onset. The primary analysis window focused on the visual epoch from 100 to 400ms after the display onset. During the specified analysis period, the average firing rate was calculated separately for TP and TA trials. To assess the ability to discriminate between TP and TA trials, the area under the receiver operating characteristic curve (AUC) was calculated (termed the target AUC). The statistical significance of the AUC was determined using the Wilcoxon rank sum test for each neuron. A cross-validation technique was used to identify the preferred target for each neuron (Ghazizadeh et al., 2018a).

### Local field potential (LFP) Analysis

To analyze the local field potential (LFP) data, the data was down-sampled to 1 kHz and subsequently subjected to a band-pass filter to retain frequencies within the range of 0.1 to 250 Hz. Then, notch filters were applied to remove power line harmonics noise. To enhance data quality, trials with LFP amplitudes that deviated beyond 3 standard deviations from the median were determined as noisy and removed from further analysis. These preprocessing steps were carried out for each session via EEGLAB (-v2022.0). A continuous wavelet transform (CWT) was used to average across all frequencies of relevance at each time point to examine the time-frequency data. An analytical Morse wavelet was used to generate the CWT (Aguiar-Conraria and Soares, 2011). Four unique frequency bands used in this study included: alpha (8-12 Hz), beta (12-30 Hz), low-gamma (30-60 Hz), and high-gamma (60-200 Hz) ranges.

### Second-order statistics of firing rate

The variability in firing rate can be separated into two distinct components: the first component pertains to the trial-to-trial firing rate variability (referred to as nRV), while the second component is associated with the within-trial spiking irregularity (referred to as nSI). We have used the method developed in Fayaz et al. (2022) to parse these components with open access code which can be found at: [https://www.ghazizadehlab.org/index.php/resources/].

### Onset detection procedure

Custom-written MATLAB functions (Ghazizadeh et al., 2018a; Ghazizadeh and Hikosaka, 2021, 2022b) were used to identify neurons that were responsive to visual stimuli. In brief, the PSTH representing the average firing rate of each neuron, aligned with the onset of the visual display, was computed for all trials. Then, the calculated firing rate was transformed into z-scores by comparing it to the baseline activity recorded during the period from -120 to 30ms relative to the display onset. The MATLAB function “@*findpeaks*” was used to identify the initial peak response that occurred after the display onset, with a minimum peak height threshold of 1.64, corresponding to the 95% confidence interval. The visual onset of the neuron was determined as the first valley observed before the first detected peak. The identical algorithm was employed to detect the initiation of the target signal, using the average PSTH of a specific neuron for target computation.

### Object-value training task

Monkeys were trained on object values using an object saccade task (Ghazizadeh and Hikosaka, 2021, 2022b), as illustrated in Fig 1B. During each trial of the task, a fixation point appeared at the center of the screen, and the subjects were instructed to maintain fixation on it. An object with a high or low value, representing either something good or bad, was presented on the screen at a location in the periphery when fixation was maintained for 200ms. After central fixation on a white dot, one object appeared on the screen at one of the eight peripheral locations (eccentricity 9.5°). After an overlap period of 400 ms, the fixation dot disappeared and the animal was required to make a saccade to the fractal. After 500 ± 100 ms of fixating the fractal, a large or small reward was delivered (biased reward training). After the reward was given, a varying time inter-trial interval (ITI) lasting from 1 to 1.5s began, during which a blank screen was displayed. A value training set was done with a set of 8 fractals, half of which were associated with low reward and the other half with high reward (referred to as bad and good fractals, respectively). In each value training set, there were a total of 80 trials, which consisted of 16 trials (choice) where both a good object and a bad object were simultaneously displayed on the screen, as well as 64 trials (force) where each fractal was shown 8 times in a pseudo-randomized manner. The timing arrangement for both the choice trials and the force trials was identical, with the only difference being that in the choice trials, the monkey had to make a saccade to choose one of the objects to receive its corresponding reward. When the monkey completed a trial, a correct auditory voice was presented, while an error auditory voice was presented in the case of a break in fixation or an early saccade. The monkey’s object value knowledge was evaluated using the results of the choice trials.

### Value-driven search task

Subjects had to choose an object with high value (good) among a varying number of objects with low value (bad) in target-present (TP) trials or reject the trials by staying at or going back to the center in target-absent (TA) trials when every object exhibited was bad (Ghazizadeh et al., 2016). Before the search task, monkeys completed a separate object value training task where they acquired knowledge regarding the object-value associations. Target-present (TP) and target-absent (TA) trials were randomly mixed with equal probability. The task started with the appearance of a purple fixation dot (Fig 2E). After 400 ms of fixation, a display with 3, 5, 7, or 9 fractals was turned on and the fixation point was turned off. Fractals were arranged equidistant from each other on an imaginary 9.5° radius circle. The location of the first fractal (arbitrary) was uniformly distributed around this circle. Each block of the search task consisted of 240 correct trials. For each trial, the fractals were selected in a pseudo-random manner from a set of 24 fractals, which included 12 “good” and 12 “bad” fractals. This set also included three sets of 8 fractals each (4 good and 4 bad), which the monkeys learned either >5 days or in 1-day in an object value training task. The monkey had a time of 3s to either select an object by fixating on it for 600ms (commitment time) and reject the trial by returning gaze to the center. Upon choosing an object, the reward associated with the selected object was given and the display was turned off. An ITI of 1-1.5s followed after receiving the reward. If the monkey’s gaze was interrupted within a 100ms timeframe after the committing time, an error tone would be produced. However, the monkey had the liberty to shift its gaze away from an object before the commitment time had lapsed. A trial could be rejected in two ways: by maintaining gaze on the center dot for 900ms after the display onset or by returning to the center and fixating on the fixation dot for 300ms. Following the rejection, the next trial promptly commenced after a delay of 400ms, and there was a 50% chance that the subsequent trial would feature a good object. In addition, monkeys had a 40% chance of obtaining a medium reward (60% of the larger reward) for the rejection trials. The purpose of this was to encourage the animals to reject TA trials if they happened to occur in a sequence and discourage them from choosing bad objects. Monkeys typically finished 240 correct trials in each search block. Subjects were well-trained in the search task prior to this experiment (>10 search practice sessions).

**Fig 2.**
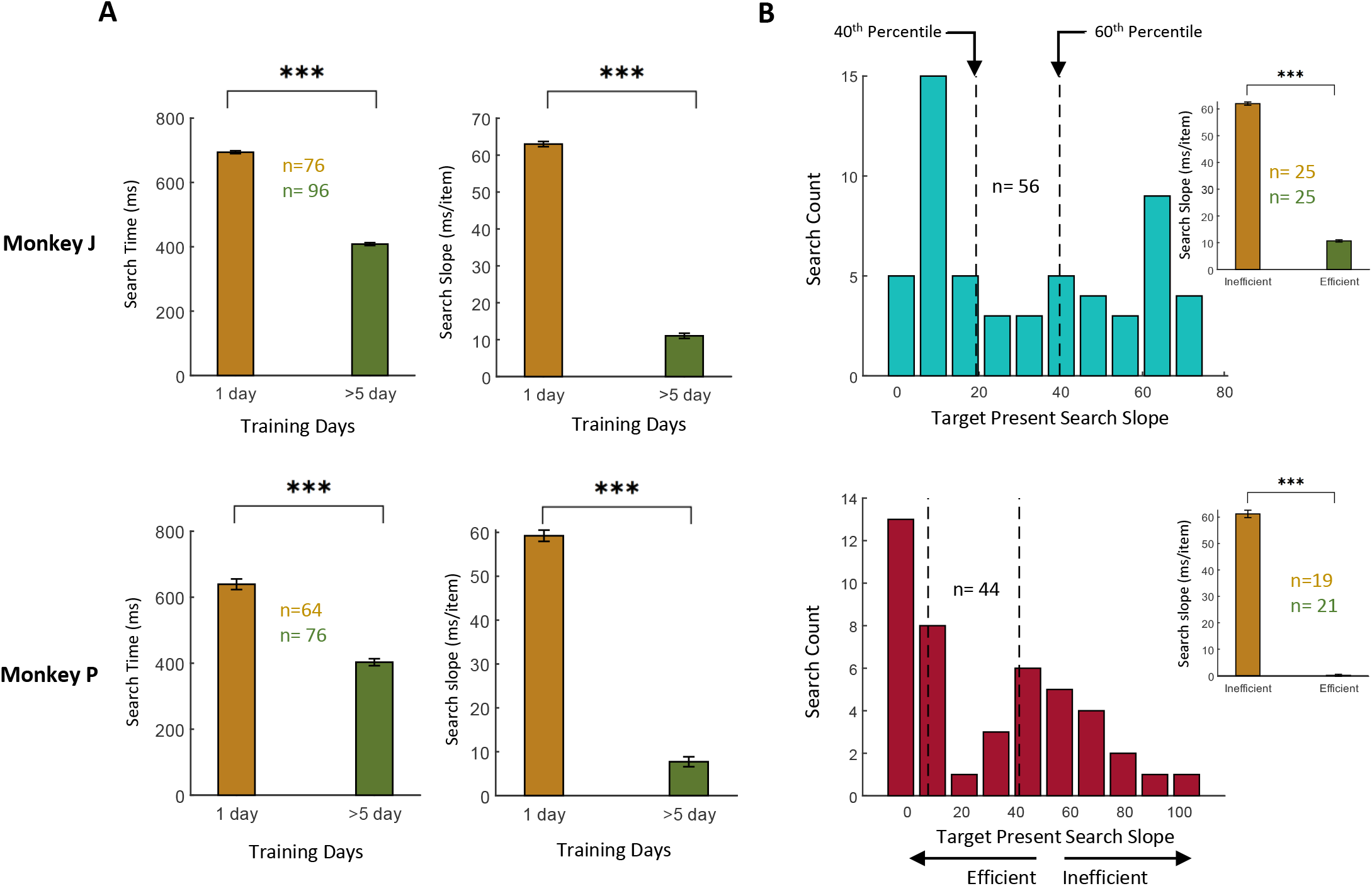
Behavioral performance of monkeys in value-based visual search. A) Search time and search time slope, respectively, for one-day trained and more than five-day trained fractal sets in TP trials for Monkey J (top) and Monkey P (bottom). B) Search time slope distribution for Monkey J (top) and Monkey P (bottom) in TP trials for all sessions. The x-axis shows the search time slope, and the y-axis indicates the number of sessions. Sessions with search slopes in the lower 40th percentile or higher 60th percentile were grouped as efficient and inefficient, respectively, for each monkey for subsequent analysis. Insets show the average search time slope for inefficient (orange) and efficient (green) search sessions for each monkey based on the percentile definition. The symbols *, **, and *** indicate statistical significance levels at p < 0.05, p < 0.01, and p < 0.001, respectively.

### One-search and two-search neurons

Each neuron was recorded with either one or two search blocks with 1-day or >5-day value trained fractals. Neurons that only had a single search block were categorized as “single-search neurons,” whereas those recorded during two search blocks were labeled as “two-search neurons”. In the two-search neurons, one search block involved the use of fractal sets that were over-trained, while the other block utilized fractal sets that were 1-day trained in a counter-balanced fashion.

### Efficient and inefficient search types

The efficiency of the search sessions was determined based on the search slope during the TP trials (Wolfe, 2000). For each session, the search slope was calculated using MATLAB’s *“@regress*” function. We calculated the average search slope in TP (target present) trials for each search block separately. For neurons with two search blocks (two-search neurons), the session with the lower search slope was classified as efficient, while the other session was deemed inefficient if the difference in slope exceeded 10ms per item. For single-search neurons, recording sessions with a search slope below the 40th percentile of the corresponding monkey’s search slopes were categorized as efficient, while sessions with a search slope above the 60th percentile were considered inefficient.

### Target preference cross-validation

The target preference of each neuron was determined using a cross validation method (Ghazizadeh et al., 2018a). The sign of firing difference between TP and TA trials in odd trials was used to determine target preference in even trials and vise-versa (i.e., if a neuron fired higher for TA compared to TP in odd trials, this firing preference label was applied to even trials and similarly target preference label in odd trials came from even trials). The odd and even trials were then combined using their cross validated target preference labels. Average firing to preferred and non-preferred target and AUC of preferred versus non preferred target were constructed using these cross validated target preferences for each neuron. All neurons were used in AUC averages whether target preferences were significant or not. To determine the TP and TA discriminability in SNr, the area under the receiver operating characteristic curve was calculated for each neuron (target AUC). Target AUCs above 0.5 indicate higher firing in TP compared to TA trials or TP-preference, and below 0.5 indicates higher firing in TA compared to TP trials or TA-preference (Fig 4G).

### Utilizing Supervised Learning with SVM Classifier for Investigating Neural Correlates of Search Efficiency

We utilized a supervised learning approach with a support vector machine (SVM) classifier. To train the classifier, we split the dataset into training and validation sets. In our study, we employed an 80/20 split, where 80% of the data were used for training and 20% for validation. In this study, each search block was categorized into either an efficient or inefficient search type (ground truth) based on the TP search slopes, as described before. Subsequently, a SVM was trained to predict the search type by utilizing the average target signal between 100-400ms after the display onset in each search block. The classification accuracy in the validation data was then reported.

### Statistical tests and significance levels

A one-way ANOVA test was conducted to determine the significance of set size on search time, accuracy, and target signal. Additionally, a two-way ANOVA test was employed to determine the effects of search efficiency and set size on the same variables. Neural firing rates were compared using t-tests for TP and TA trials in efficient and inefficient search conditions. The significance level for all statistical analyses in this study was set at P < 0.05.

## Results

In order to understand the role of the SNr in facilitating efficient search, responses of SNr neurons were recorded while monkeys engaged in a visual search task for a high-value fractal object (methods section for more information). The value of fractals was learned using an object-value training task prior to the search task (value training, see Fig 1A-B). A value training session was done with a set of 8 fractals, half of which were associated with low reward and the other half with high reward (referred to as bad and good fractals, respectively). The majority of trials (80%) during value learning consisted of force trials where a single fractal was shown in the periphery and after the fixation offset, the monkey had to saccade to and fixate the fractal to receive its associated reward. Choice trials (20%) interspersed during the value training were used to assess the subject’s knowledge of fractal values and consisted of two diametrically presented fractals between which monkeys had to choose by making a saccade and receiving the corresponding reward (Fig 1B). It was previously shown that value-based visual search can be inefficient or efficient depending on the duration of value training that preceded search (Ghazizadeh et al., 2016; Abbaszadeh et al., 2023). Thus, to create both efficient and inefficient search, the object sets underwent training for either 1-day or >5 days (over-trained fractals).

Acquisition times and choice rates during the value training sessions show robust value learning in a value learning session (Fig 1C). In both 1-day and 5-day groups the target acquisition times for good objects were significantly faster than for bad objects (t_220_ = 11.98, P < 8.7381e-26), consistent with a higher motivation to receive the large reward associated with good objects. Furthermore, the choice rate in the 1-day and 5-day groups were high and significantly above change averaging 92% and 100%, respectively. These results show that animals had a very good knowledge of fractal values even after 1 day of value training with new fractals. Following last reward training, 1-day and 5-day fractals were used in visual search task during which neuronal responses in SNr were recorded (Fig 1D).

### Rewards over-training leads to efficient search

In the search task, the monkeys were required to locate a single good fractal among multiple bad fractals in target-present (TP) trials to receive high reward. In target-absent (TA) trials where all the displayed objects were bad, they had to reject the trial by returning to the center. Rejection of a TA trial, resulted in a rapid progression to the next trial while choosing any of the bad objects was followed by the delivery of a small reward and the normal ITI to the next trial (as shown in Fig 1E). During the TP trials, monkeys received a large reward for finding the good fractal and a small reward for choosing any of the bad fractals. While fractal values in both groups were well-learned (Fig 1C), both search times and search slopes (per item of set size) were significantly smaller for overtrained fractals compared to those trained for only one day during target-present trials in both monkeys (Fig 2A). The monkeys’ behavioral search slopes varied from serial (with slopes greater than 40ms/item) to parallel (with slopes close to zero ms/item) during TP trials (Fig 2B). We grouped search performance into two types based on search slopes in TP trials (see methods).

The accuracy of good object choice in TP trials was significantly higher in efficient vs inefficient search across all set sizes (90% vs 65%, efficiency main effect: F_1, 372_ = 223.53, p<0.001, Fig 1F). Similarly, the accuracy of trial rejection in TA trials was significantly higher in efficient vs inefficient search (Efficiency main effect: F_1, 372_ = 85.20, p<0.001). The accuracy in TP was not set -size dependent (set size main effect: F_3, 372_ =0.31, p=.81, interaction: F_3,372_ = 0.92, p=0.42, Fig 1F) but there was a significant reduction of accuracy in both groups in TA trials in higher set sizes (set size main effect: F_3, 372_ = 9.23, p<0.001, interaction: F_3, 372_ =0.06, p=0.97, Fig 1F).

Search time was significantly higher in inefficient vs efficient search in both TP (F_1,372_ = 809.05, p<0.001, Fig 1F) and TA trials (F_1,372_ = 29.93, p<0.001, Fig 1F). There was main effect in set size in both TP (F_3,372_ = 87.98, p<0.001) and TA trials (F_3,372,_ = 60.78, p<0.001). As expected from our search type definition there was a significant interaction between search type and set size in TP search slopes (interaction: F_3,372_ = 46.09, p<0.001). Additionally, there was a trending interaction effect between search type and set size in TA trials (interaction: F_3,372_ = 2.27, p=0.08). Indeed, in TA trials there was a significant reduction of search slopes from ∼90ms/item in inefficient searches to ∼55ms/item in efficient searches (Fig 1F).

### SNr differential response to TP and TA trials is enhanced in efficient search

To investigate the role of SNr in value-based search efficiency, the activity of single neurons was recorded in SNr region (Fig 1G) and contrasted in efficient and inefficient search types. Fig 3 shows the example response of SNr neuron in both search types in TA and TP trials and separately for each set size with the firing rasters. As can be seen the neuron is inhibited in both TP and TA trials in inefficient search but the inhibition is stronger in TP trials. During efficient search the inhibition to TP trials deepens while TA response shows a late excitation, there was a marked inhibition in firing in TP trials. Fig 4A shows the average population response which shows similar firing dynamics as the example neurons shown. As can be seen in the efficient search, there was a marked inhibition in firing in TP trials. There was also a set-size dependance in firing, such that there was a larger inhibition in smaller set-sizes. This set-size dependence was significantly larger in inefficient compared to efficient search consistent with the larger effect of set-size on behavior in inefficient search (Fig 4B). While SNr neurons showed inhibition in inefficient search during TA trials, there was a slight excitation in TA trials in efficient search. Once more the set-size dependence consisted of higher firing to larger set-sizes and was significantly larger in inefficient compared to efficient search (Fig 4B). The inhibition in TP and excitation in TA trials in efficient search resembled responses previously reported for a single good or bad object, respectively (Yasuda et al., 2012; Ghazizadeh and Hikosaka, 2022b). Notably, during inefficient search SNr showed an inhibition in both TP and TA trials consistent with the difficulty that the animal faced in determining the absence of a good object in the display.

**Fig 3.**
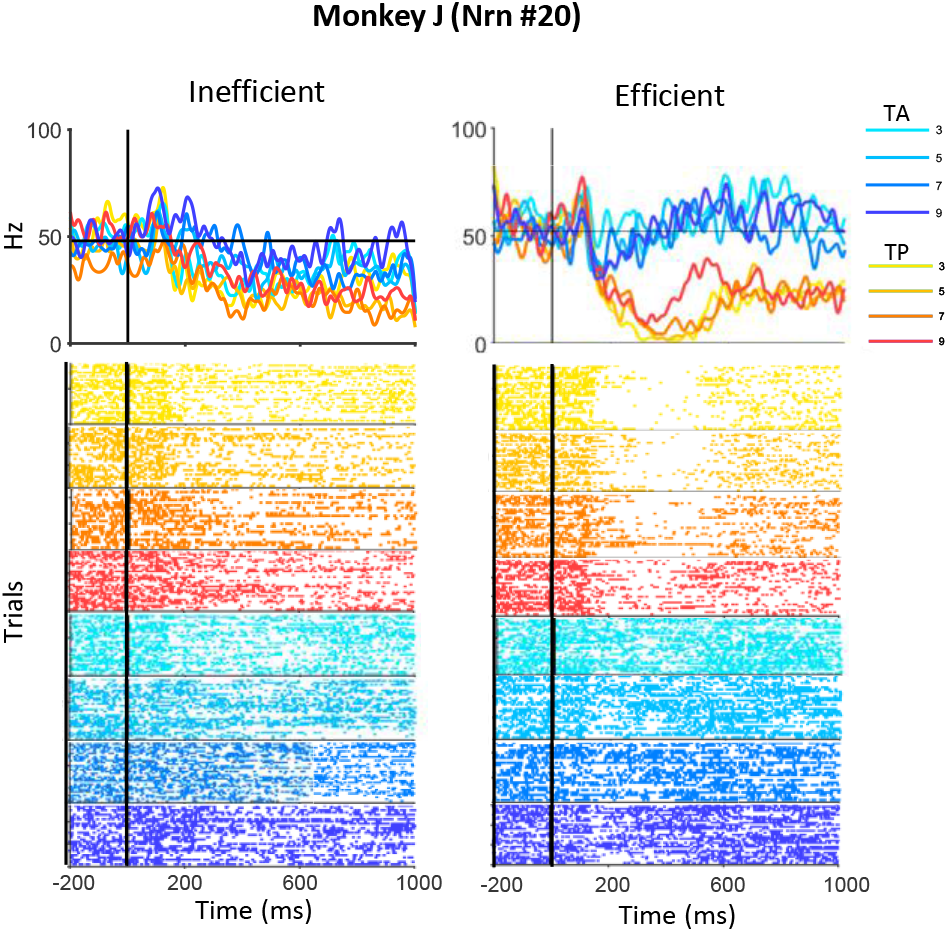
Firing rate response to target-present (TP) and target-absent (TA) trials during inefficient and efficient searches in an example SNr neuron. Peristimulus time histograms (PSTH) and raster plots in inefficient (left) and efficient (right) searches, time-locked to display onset are shown. The set sizes of 3, 5, 7, and 9 are shown in red gradient colors for TP trials and blue gradient colors for TA trials, and inefficient and efficient search types are shown on the left and right sides. The mean firing rate for the baseline period (-120 to 30ms) is indicated by a black horizontal line.

The firing rate in TA trials was significantly higher in efficient vs inefficient search across all set-sizes (main effect: F_1, 372_ = 6.60, p<0.05, Fig 4C). Additionally, there was a tendency for the firing rate in TP trials to be higher during inefficient search compared to efficient search (main effect: F_1, 372_ = 3.47, p=0.06, Fig 4C).

**Fig 4.**
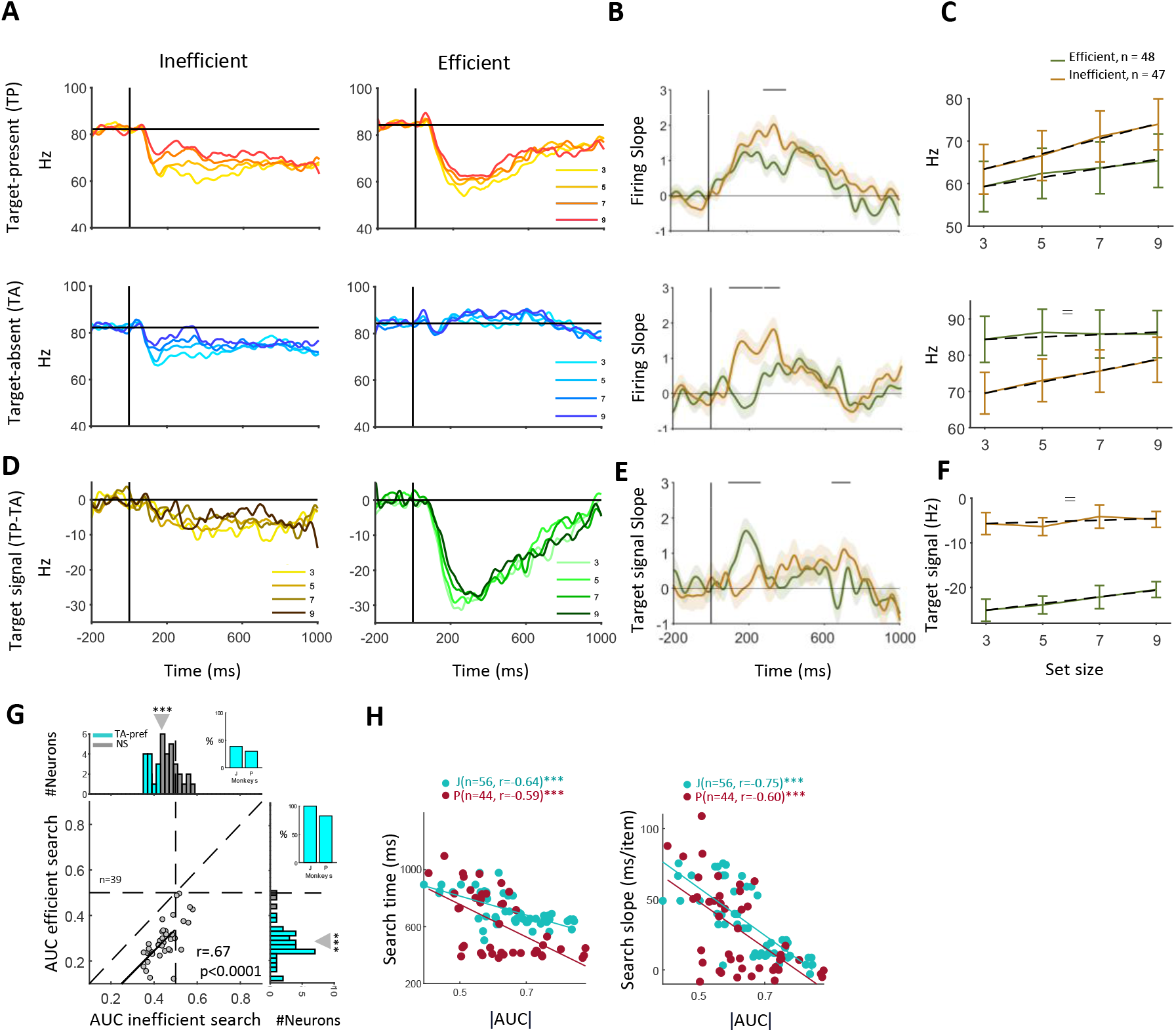
The SNr differential neural firing in TP and TA trials correlates with search efficiency. A) The population average firing time-locked to display onset separately for set sizes in TP and TA trials. The mean firing rate for the baseline period (-120 to 30ms) is indicated by a black horizontal line. B) The firing slope as a function of set size in inefficient (left side) vs. efficient (right side) searches in TP (top row) and TA (bottom row) trials. Lines above the PSTHs indicate significance at the p < 0.01 in 25ms bins. C) The average firing rate across neurons during visual epoch (100-400ms relative to display onset) across the set size for both the inefficient (orange) and the efficient (green) searches for TP (top) and TA (bottom) trials. The dashed lines show the linear fit of the firing rate as a function of set size. D) The firing difference between TP and TA signals (firing target signal), separately for each set size. E) Same format as B but for target signal. F) Same format as C but for target signal slope as a function of set size. G) Target signal AUC scatter for neurons in inefficient vs efficient search. The linear fit is shown as a solid dark line. Marginal target signal AUC histograms are shown and neurons with significant target signal AUC are color-coded (TA-preferred, significant AUC < 0.5, and nonsignificant NS neurons). The insets show the percentage of TA-preference neurons for both monkeys. H) Scatter plots showing search time and search slopes in TP trials as a function of the absolute target signal AUC in firing rate across sessions for the two monkeys. The symbols =, /, and x represent the main effects of search efficiency, set size, and their interaction, respectively.

The target signal was negative in SNr consistent with the response to single objects (Yasuda et al., 2012) (Fig 4D). Importantly, the size of the target signal was much larger in efficient compared to inefficient search in visual epoch (t_93_=-7.56, p<0.001, Fig 4E). The main effect of the search type on the target signal was significant for the visual epoch (F_1, 372_ = 160.29, p<0.001, Fig 4F). Compared to TP or TA trial, the set-size dependence in the target signal (TP minus TA firing) was less prominent. The efficient search target signal showed higher set-size dependence compared to inefficient search early in the trial (∼200ms) while later in the trial (600-800ms) the target signal in inefficient search had higher set-size dependence (Fig 4D). The onset of the target signal across the population was 105±8.27ms after display onset which is rapid and comparable to the onset of the target signal when a single object is shown (98.92±6.25ms) (Ghazizadeh and Hikosaka, 2021).

To determine the TP and TA discriminability in SNr, the area under the receiver operating characteristic curve was calculated for each neuron (target AUC). Value AUCs above 0.5 indicate higher firing in TP compared to TA trials or TP-preference, and below 0.5 indicates higher firing in TA compared to TP trials or TA-preference. In both search types, the average target AUC across the population was significantly below 0.5 in neurons with both search types (Fig 4G, Efficient: AUC = 0.28, t_38_ = -15.10, P < 1.2039e-17, Inefficient: AUC = 0.44, t_38_ =-5.38, P < 3.1257e-08), revealing a significant TA-preference across the SNr population (X^2^ = 72.15, p < 0.0001, Fig 4G) . TA preference was found in 100.00% and 85.00% for Monkey J and Monkey P, respectively in efficient search while in inefficient search this number was 31.81% and 36.84% for Monkey J and Monkey P, respectively for two search neurons, as can be seen in Fig 4G (for all neurons: 100.00% and 81.25% for Monkey J and Monkey P, respectively in efficient search; 35.29% and 40.00% for Monkey J and Monkey P, respectively in inefficient search). Previous results showed that only a small minority of SNr neurons had TP preference (Yasuda et al., 2012; Ghazizadeh and Hikosaka, 2021). The fact that we do not find any SNr neuron with significant TP-preference could be due to task differences as previous work was mostly done with passive viewing task while the current task is an active search task.

Importantly, plotting the target signal in efficient vs inefficient search across neurons with both search types, showed that there was a significant correlation between target AUCs in the two search types for a given neuron (r = 0.67, p < 0.0001, Fig 4G). This meant that the neurons with a stronger target signal in one search type tended to have a stronger target signal in the other search type. Additionally, the target signal in the efficient search was stronger than the inefficient search (below the unity line). Indeed, a classifier trained on the target signals of two-search neurons could discriminate the two search types with 80.8% accuracy (see methods).

Critically, results also showed a significant correlation between the average target signal of the recorded neuron in the visual epoch (100-400ms post display onset) in each session and search efficiency in that session as measured by both search time and search slope in both monkeys (monkey J, search slope: r=-0.75, p < 0.0001, search time: r=-0.60, p < 0.0001, monkey P, search slope: r=-0.64, p < 0.0001, search time: r=-0.59, p < 0.0001, Fig 4H). Therefore, one concludes that larger target signal in SNr was concurrent with faster and more efficient value-driven visual search.

Fig 5 shows the average population response in inefficient (top) and efficient (bottom) collapsed over set sizes time locked to display onset (Fig 5A) which continued to the time of saccade (Fig 5B). The difference in firing to TP and TA trials in a given set size can arise from sensory component which indicates presence of a single good object or could reflect the consequent motoric difference that results from saccading to the target. In this study, we refer to this overall difference in firing as the target signal, which potentially encompasses both the sensory processing and motor differences between TP and TA trials. We note however that our preliminary analysis does not show a component of firing that is time-locked to the saccade onset (Fig 5B). Furthermore, previous work has shown that SNr neurons show strong firing differences even in passive viewing when there is no saccade (Yasuda et al., 2012; Ghazizadeh and Hikosaka, 2021). Thus, one can conclude that there is evidence in favor of sensory processing difference to be a key component in the target signal in SNr.

**Fig 5.**
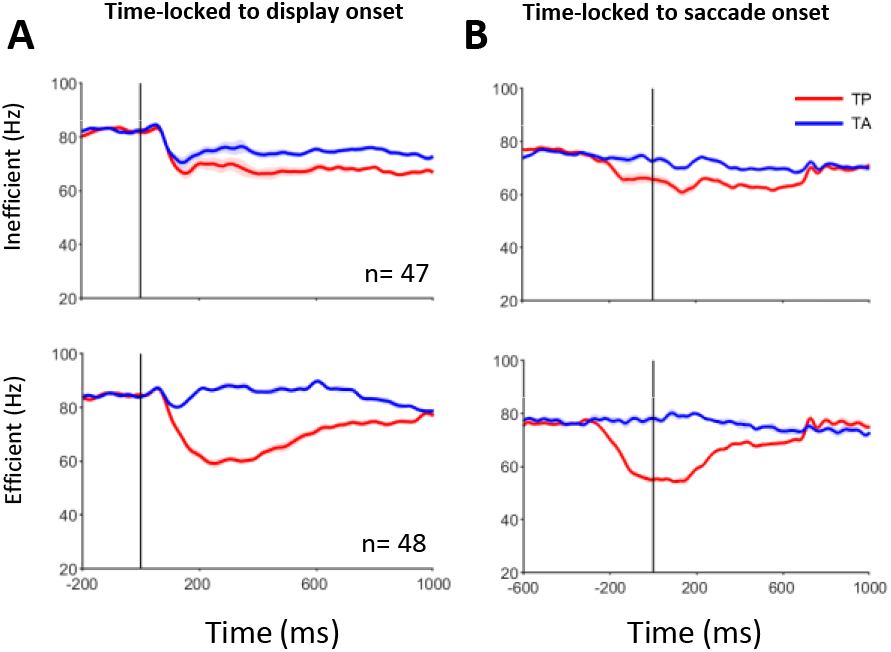
The SNr neurons in inefficient and efficient search. A) Population average PSTH time locked to display onset in the inefficient (top) and efficient (bottom) searches. B) Same format as A, but time-locked to saccade onset.

### SNr exhibits higher spiking variability during efficient search in TP trials

Apart from the relationship between search efficiency and SNr mean firing rate (first-order statistics), we have also looked at the changes in firing variability in SNr across the two search types (second-order statistics). The variability in firing rate can be separated into two distinct components: the first component pertains to the trial-to-trial firing rate variability (referred to as nRV), while the second component is associated with the within-trial spiking irregularity (referred to as nSI) (Churchland et al., 2011; Churchland and Abbott, 2012; Fayaz et al., 2022).

Results showed a decrease in nRV following the display onset, which was followed by a subsequent increase in nRV later in the trial consistent with earlier studies (Churchland et al., 2010; Churchland et al., 2011; Fayaz et al., 2022). There was no significant difference in nRV during this epoch between efficient and inefficient searches (ps > 0.05, Fig 6A-B). On the other hand, spiking irregularity showed a marked increase following display onset. Notably, spiking irregularity was significantly higher during efficient search in TP vs TA trials (F_1,372_ = 11.00, p < 0.001, Fig 6A). The target signal in nSI (i.e. nSI difference in TP vs TA trials) was significantly larger in efficient vs inefficient search in visual epoch (t_93_ = 2.71, P < 0.01, Fig 6B) but showed significant reduction in higher set-sizes for efficient search (Fig 6C). The changes in nSI and nRV in TP and TA trials were not set-size dependent in general barring some fluctuations in time (ps > 0.05).

**Fig 6.**
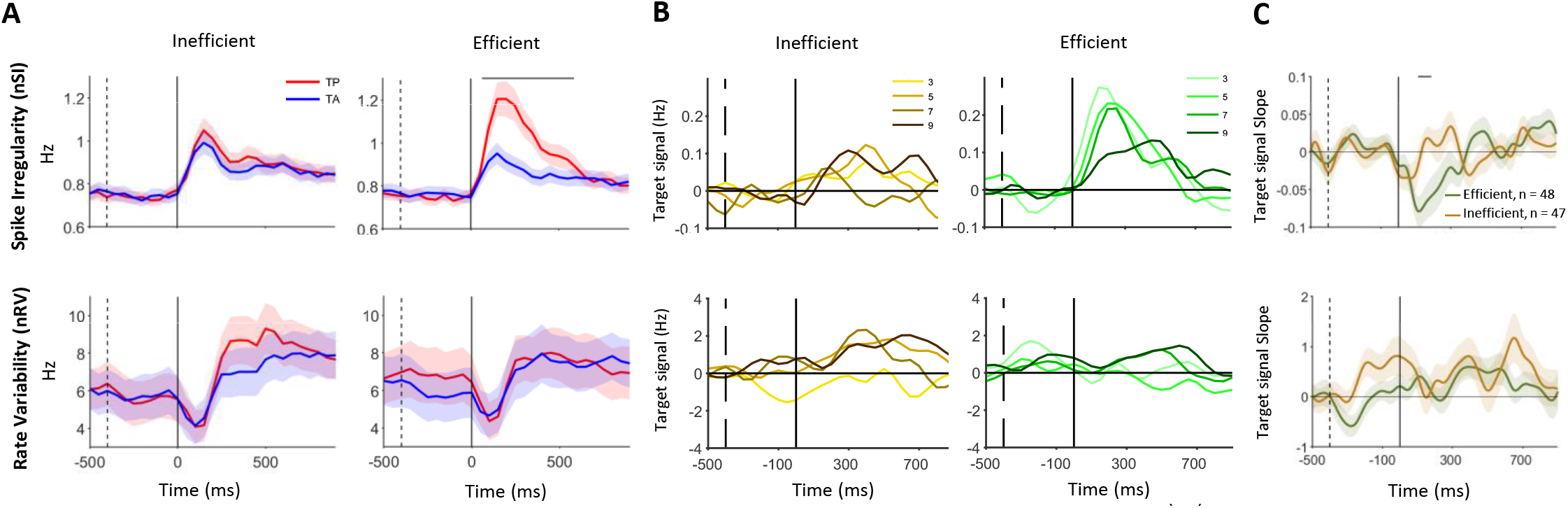
Spiking variability in TP trials during efficient searches. A) Population average collapsed across all set sizes in the TP trials (red) and the TA trials (blue) for the inefficient (left) and efficient (right) searches for spiking variability (nSI, top), and rate variability (nRV, bottom). B) The difference between TP and TA trials separately for set sizes for inefficient (left) and efficient (right) searches for nSI (top), and nRV (bottom). C) The target signal slope as function of set size across time for nSI (top), and nRV (bottom) for efficient and inefficient searches. Lines above the PSTHs indicate significance at the p < 0.01 level.

### High gamma-band power in the SNr is associated with search efficiency

Modulations in different frequency bands of the local field potential (LFP) are associated with complex cognitive processes (Buzsaki and Draguhn, 2004; Buzsáki and Wang, 2012). Using continuous wavelet transform (CWT), we found significant changes in various LFP frequencies within SNr during the visual search task (32 and 24 sessions in the inefficient and efficient searches, respectively). Specifically, following search display onset there was a noticeable reduction in high gamma band (100-200 Hz) power, as well as beta band (12-30 Hz) and alpha (8-12 Hz) powers (Fig 7A).

**Fig 7.**
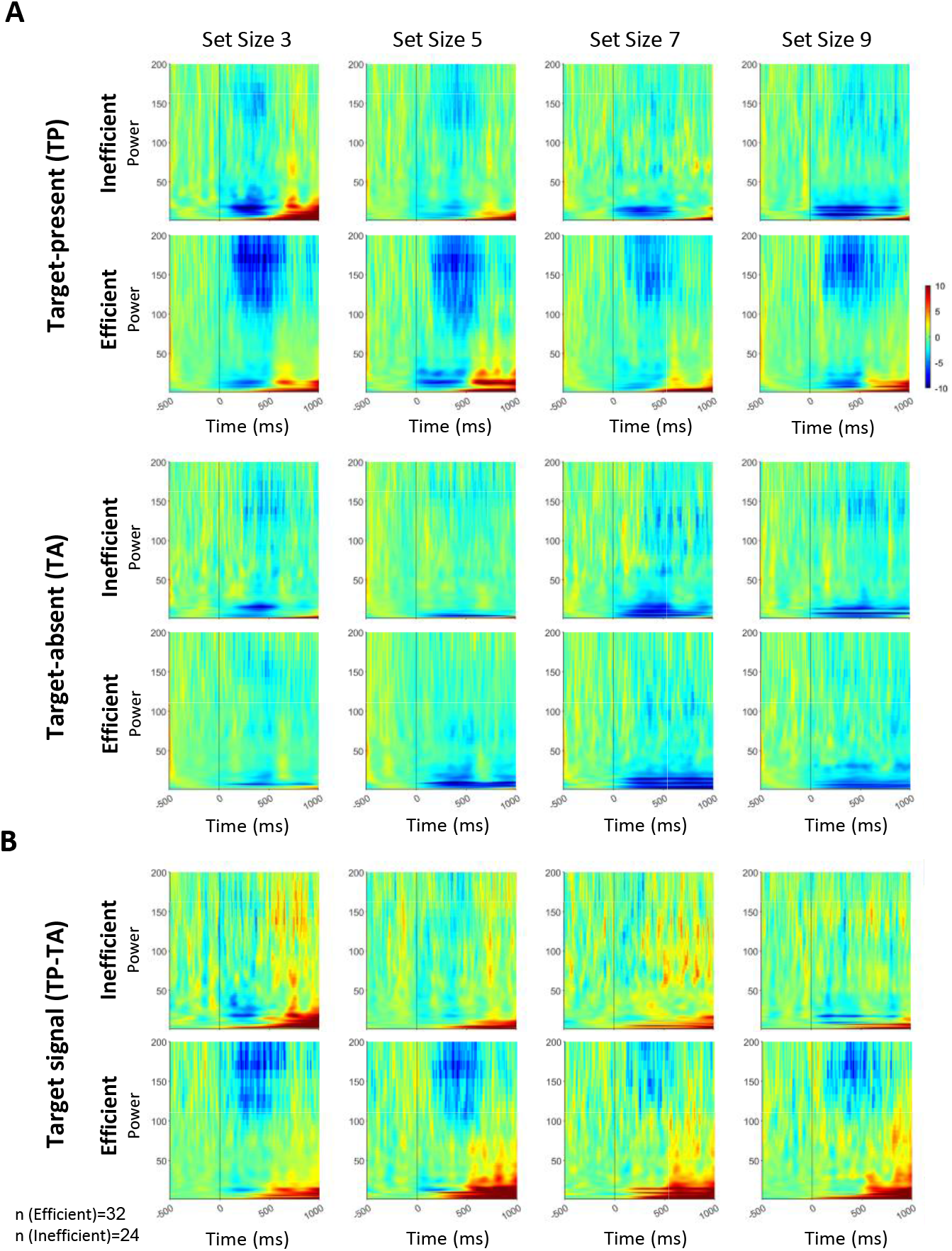
LFP power modulation time locked to display onset in inefficient and efficient searches using time-frequency analysis (CWT). A) Changes in LFP power (20log10) time-locked to display onset, spanning the frequency range of 0-200 Hz, averaged across set sizes for target-present (TP) trials (top) and target-absent (TA) trials (bottom) in inefficient and efficient search. The color code represents the changes in LFP power compared to the average baseline power (-500-0ms). B) The difference in LFP power between TP and TA trials, during inefficient and efficient search (LFP target signal).

Similar to the firing rate, one may calculate the target signal in LFP by subtracting the powers across frequencies in TP and TA trials (TP minus TA). Results showed a negative target signal in the gamma range following display onset (Fig 7B). Notably, the reduction of high gamma target signal power was significantly stronger in efficient vs. inefficient search in the visual epoch (F_1,110_ = 14.39, p < 0.0001, Fig 8), but there was no main effect in set-size or interaction between search type and set-size (Interaction: F_3,110_ = 0.66, p=0.58, set-size main effect: F_3,110_ = 0.33, p = 0.80). There was no significant difference in the target signal across alpha, beta, and low gamma bands between efficient and inefficient search in the visual epoch (ps > 0.05, Fig 8).

**Fig 8.**
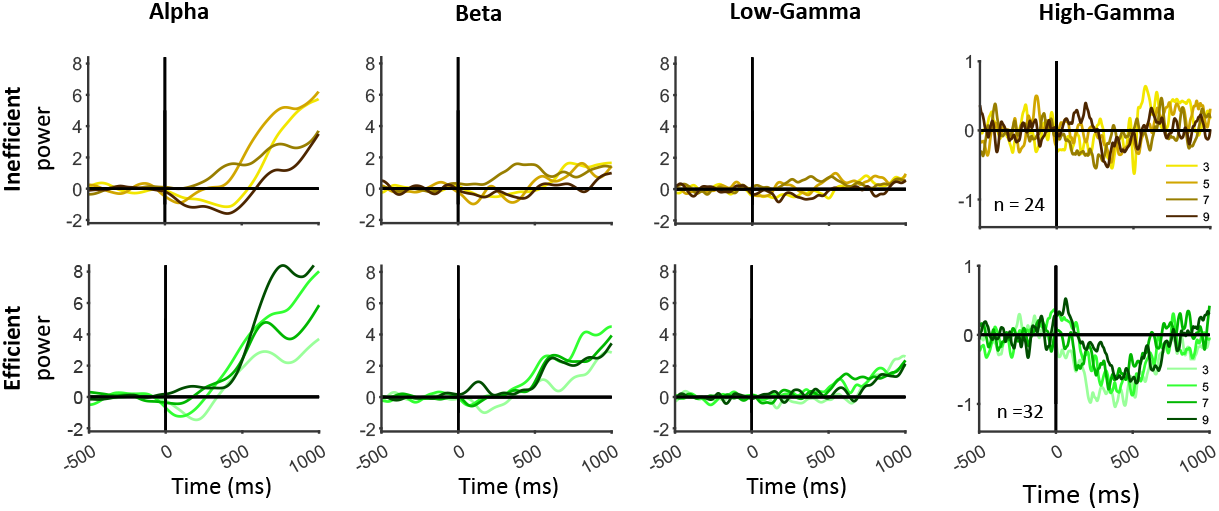
Target signals in alpha, beta, low gamma, and high-gamma LFP frequency bands during the efficient and inefficient search. Target signal in the alpha, beta, low gamma, and high-gamma frequency bands for different set sizes in LFP sessions of inefficient and efficient searches.

Similar to the target signal in firing rate, there was a significant negative correlation between the target signal during the visual epoch in high gamma and search slope in both monkeys and a trend for negative correlation with search time (monkey J, r=-0.54, p < 0.01; monkey P: r=-0.42, p < 0.05, Fig 9). No significant relationship in the visual epoch was observed in alpha, beta, or low gamma bands (except for alpha band in monkey J for search slope: r=-0.49, p < 0.01, and search time: r=-0.38, p < 0.05, Fig 9). These suggest that among frequency bands in SNr, high gamma signal encodes target signal and is a good correlate of search efficiency in the visual epoch. While the alpha band power in a later epoch (500-800ms) also showed a significant correlation with search efficiency given the fact that mean saccade times in the search task are ∼700ms this correlation would be too late to causally affect the search time (Fig 10).

**Fig 9.**
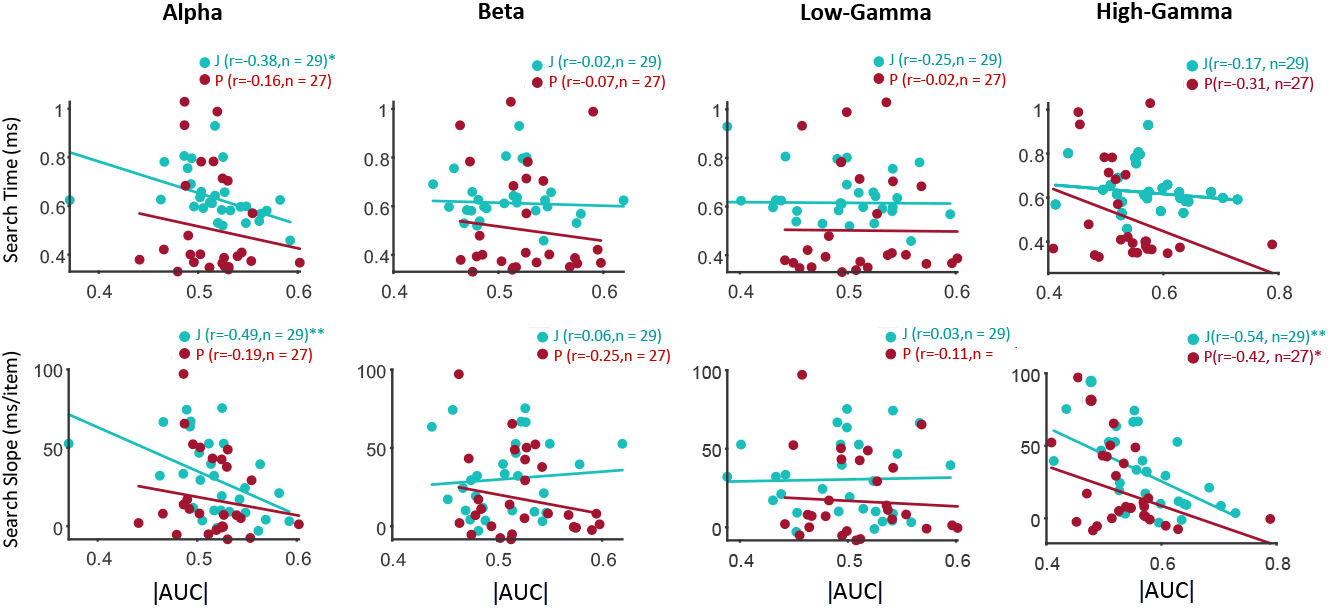
The correlation between search efficiency and target signal in alpha, beta, and low gamma bands in visual epoch (100-400ms) after display onset. Scatter plots showing search time and search slopes in TP trials as a function of the absolute target signal AUC in alpha, beta, low gamma, and high-gamma LFP frequency bands across sessions for the two monkeys.

**Fig 10.**
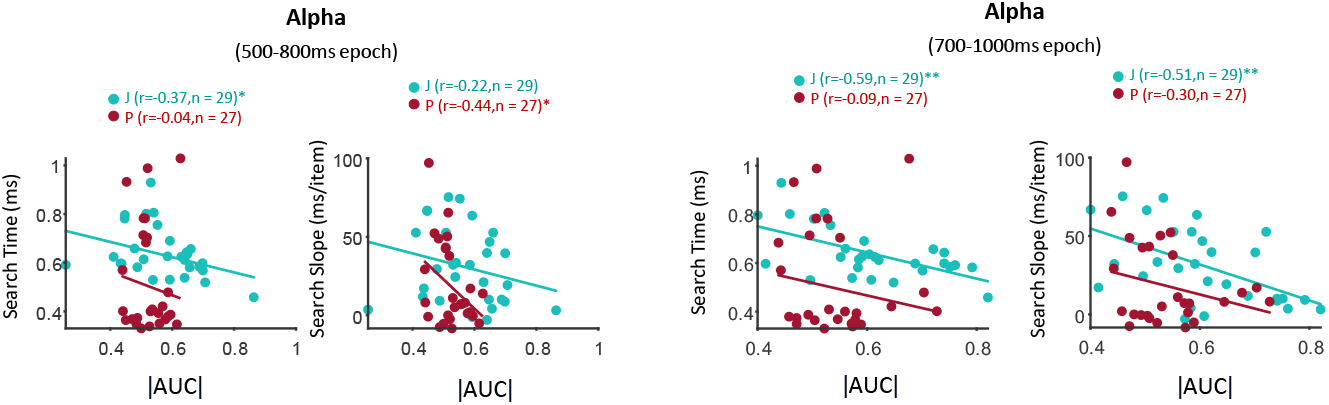
The correlation between search efficiency and target signal in alpha band in later periods after display onset. Similar to Fig 9 for alpha target signal but for 500-800ms and 700-1000ms epochs.

## Discussion

While the role of overtrained value as a high-level guiding feature in visual search has been recently recognized (Ghazizadeh et al., 2016; Wolfe, 2020), its underlying neural mechanism remains unknown. Given the role of SNr in control of visual attention and saccades (Hikosaka et al., 2006) as well as in object value memory (Yasuda et al., 2012; Ghazizadeh and Hikosaka, 2021, 2022b), we investigated its role in value-based visual search efficiency by recording neural activity of a single neuron in both efficient and inefficient search with varying set sizes. The neurons in the SNr exhibited stronger inhibition during TP (target present) trials in comparison to TA (target absent) trials (Fig 3, Fig 4A). Results show that the firing rate in TP and TA trials in inefficient search was more set-size dependent. Importantly, the firing difference to TP and TA trials (firing target signal) was significantly stronger in efficient search and there was a significant correlation between the strength of this target signal and search efficiency across sessions (Fig 4). In addition to the firing rate, in efficient visual search was concurrent with higher spiking irregularity following the display onset during TP trials (Fig 6). LFP signals in SNr were also affected by search efficiency. In particular, high gamma power showed a larger reduction in TP trials during efficient search and this power modulation was correlated with search efficiency in a given session (Fig 9).

Efficient search for valuable objects is vital for survival in the wild. Animals explore their environment, encountering objects of varying values. Objects are considered high value and preferred, based on their past learned associations with reward and their effect on enhancing the animal’s competitive fitness. Objects associated with rewards tend to capture gaze and attention independent of low-level salience (Libera and Chelazzi, 2006; Hickey et al., 2010; Anderson et al., 2011; Towal et al., 2013). Our results showed that object value can also result in efficient object search (Ghazizadeh et al., 2016). Recent evidence shows that areas within BG (posterior BG circuitry) play a key role in formation and retention of object value memories and in directing gaze toward valuable objects (Yamamoto et al., 2012; Yasuda et al., 2012; Kim and Hikosaka, 2013; Kim et al., 2017; Kunimatsu et al., 2019) modulated by inputs from a special class of dopaminergic neurons in the SNc (Kim et al., 2015). The output of BG emanating from SNr impacts gaze direction via projection to the SC (Hikosaka and Wurtz, 1983; Hikosaka et al., 2019). Target signals in the posterior BG circuit can also affect cortical regions through thalamocortical connections, facilitating quick identification of valuable objects (Middleton and Strick, 2002; Ghazizadeh et al., 2018b). Given the inhibitory role of SNr on saccades, the stronger reduction of its neural firing in TP trials during efficient search should result in greater disinhibition of SC neurons to allow rapid gaze shift toward detected targets (Basso and Wurtz, 1997).

We have also looked at the changes in the neural variability using a recent method that allows parsing two components related to firing rate variability (nRV) and spiking irregularity (nSI) (Vinci et al., 2016; Fayaz et al., 2022). The temporal dynamics of nRV and nSI in SNr in the search task was similar to its pattern in passive viewing of good and bad objects (Fayaz et al., 2022). Consistent with previous report (Churchland et al., 2010), we found a transient reduction of firing rate variability (nRV) following display onset but unlike the finding in some cortical areas (Churchland et al., 2011), this component of variability was not set-size dependent. The nRV was not modulated by search efficiency either. Unlike nRV, spiking irregularity (nSI) was found to be affected by search efficiency. In particular, efficient search was associated with higher spiking irregularity in the SNr during target-present trials compared to target-absent trials. While our understanding of the computational and behavioral impact of spiking irregularity is nascent, there is growing evidence implicating spiking irregularity in information coding and for feasibility of its decoding in downstream areas (Gallinaro and Clopath, 2021) as well as its role for affecting behavior (Doron et al., 2014). In particular it is shown that bursting can contributes to higher spiking irregularity in SNr (Fayaz et al., 2022) and that bursting increases the reliability of synaptic transmission (Zeldenrust et al., 2018). It remains to be seen whether higher spiking irregularity during efficient search indeed facilitates detection and saccade to good objects via a more effective information transmission to SC for instance.

Despite a large number of studies on SNr neural firing in value-related tasks there is surprisingly little work on the LFP signals in this region especially using event-related time-frequency analysis. This study provides the first systematic analysis of LFP in a complex value-based task in SNr. Time-frequency analysis of LFP showed clear reduction of power in high gamma band (100-200 Hz), beta band (12-30 Hz) and alpha band (8-12 Hz) time-locked to display onset in both TP and TA trials. The power reduction was more prominent in TP compared to TA trials in efficient search in high gamma band. This high gamma target signal was correlated with improved search slopes across sessions (Fig 9). While the origin and significance of LFP signals in cortical and subcortical regions can be very diverse (Buzsáki et al., 2012), it is generally thought that while high gamma band modulation reflects local computations and carry significant mutual information with the spikes in a region, low frequency band codes information independent from firing rates carrying mostly afferent information to the region (Belitski et al., 2008; Rasch et al., 2008; Khawaja et al., 2009). This interpretation suggests the intriguing possibility that given the presence of target signal in high gamma and its absence in lower frequencies, rather than being fully determined by inputs to SNr, the efficient detection of high value target is computed withing SNr neurons. The enhanced target signal in high gamma band during efficient value-based search has been recently reported in ventrolateral prefrontal cortex (vlPFC) (Abbaszadeh et al., 2023). vlPFC and SNr are known to participate in cortico-basal ganglia loops and to commonly encode object value memories (Ghazizadeh and Hikosaka, 2021). Overall, these findings suggest that gamma-band activity in the SNr may play a critical role in visual search, potentially serving as a marker for target detection and attentional selection.

The current findings suggest that the disinhibition of SC by SNr can enable efficient identification of high-value objects. The results revealed a link between the strength of the target signal in the firing rate and gamma band activity in the SNr and the efficiency of visual search for objects. The relatively large size of receptive fields in SNr (Ghazizadeh and Hikosaka, 2021) allows for parallel processing of contralateral visual hemifield to indicate presence of a valuable target. It is then possible for vlPFC which is sensitive to object value and has a more localized receptive field (Ghazizadeh and Hikosaka, 2021) to direct gaze toward the exact location of the valuable target (Abbaszadeh et al., 2023). Additional research is needed to establish and differentiate the role of cortical and subcortical areas involved in ‘value pop-out’ during visual search for objects.

## Acknowledgments

We thank M. Khorasani, A. Shabani, and G. Sadeghian, for technical assistance in monkey care and Ghazizadeh lab for helpful cooperation (S. Meghdadi, A. Panjehpoor, M. Khademian, A. Mirzadeh, Nikkozadeh and Mokhtari).

## Funding

This work was financially supported by the Cognitive Science and Technology Council of Iran (CSTC) (Grant No. 14071/100/d).

## Competing interests

The authors declare that they have no competing interests.

## Data and materials availability

All data needed to evaluate the conclusions in the paper are present in the paper. Additional data related to this paper may be requested from the authors.

## Notes

### Competing Interest Statement

The authors have declared no competing interest.

### Summary of Updates

Figures updated; Manuscript file updated.

